# Heterologous expression of cryptomaldamide in a cyanobacterial host

**DOI:** 10.1101/2020.08.26.267179

**Authors:** Arnaud Taton, Andrew Ecker, Brienna Diaz, Nathan A. Moss, Brooke Anderson, Raphael Reher, Tiago F. Leão, Ryan Simkovsky, Pieter C. Dorrestein, Lena Gerwick, William H. Gerwick, James W. Golden

**Affiliations:** Division of Biological Sciences, University of California, San Diego, La Jolla, California 92093, United States; Center for Marine Biotechnology and Biomedicine, Scripps Institution of Oceanography, University of California, San Diego, La Jolla, California 92093, United States; Skaggs School of Pharmacy and Pharmaceutical Sciences, University of California, San Diego, La Jolla, California 92093, United States

**Keywords:** Cyanobacteria, natural products, heterologous expression, cryptomaldamide

## Abstract

Filamentous marine cyanobacteria make a variety of bioactive molecules that are produced by polyketide synthases, non-ribosomal peptide synthetases, and hybrid pathways that are encoded by large biosynthetic gene clusters. These cyanobacterial natural products represent potential drugs leads; however, thorough pharmacological investigations have been impeded by the limited quantity of compound that is typically available from the native organisms. Additionally, investigations of the biosynthetic gene clusters and enzymatic pathways have been difficult due to the inability to conduct genetic manipulations in the native producers. Here we report a set of genetic tools for the heterologous expression of biosynthetic gene clusters in the cyanobacteria *Synechococcus elongatus* PCC 7942 and *Anabaena* (*Nostoc*) PCC 7120. To facilitate the transfer of gene clusters in both strains, we engineered a strain of *Anabaena* that contains *S. elongatus* homologous sequences for chromosomal recombination at a neutral site and devised a CRISPR-based strategy to efficiently obtain segregated double recombinant clones of *Anabaena*. These genetic tools were used to express the large 28.7 kb cryptomaldamide biosynthetic gene cluster from the marine cyanobacterium *Moorena* (*Moorea*) *producens* JHB in both model strains. *S. elongatus* did not produce cryptomaldamide, however high-titer production of cryptomaldamide was obtained in *Anabaena*. The methods developed in this study will facilitate the heterologous expression of biosynthetic gene clusters isolated from marine cyanobacteria and complex metagenomic samples.

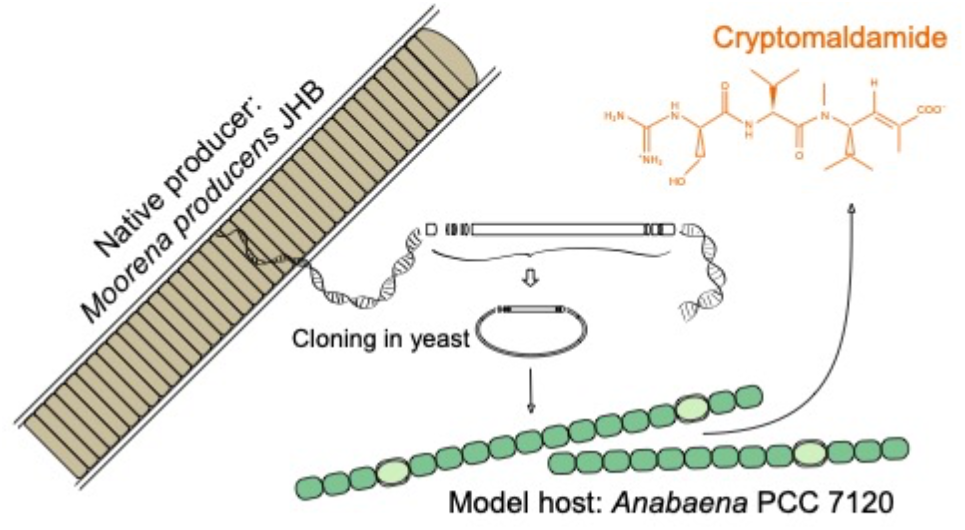

## INTRODUCTION

Cyanobacteria are sources of diverse bioactive secondary metabolites including toxins and other natural products (NPs).^1, 2^ Some species are notorious contributors to harmful algal blooms, particularly in freshwater lakes, ponds, and reservoirs, where they release various toxic molecules and sometimes cause animal and human health issues.^3^ Many cyanobacteria, whether from freshwater, terrestrial, or marine environments, carry large biosynthetic gene clusters (BGCs) that encode for the biosynthesis of diverse bioactive molecules.^4–6^ The definitive roles of these compounds in nature are still elusive, but they may serve as signaling molecules, toxins, or allelochemicals that may inhibit competitors or act as a deterrent to predators.^7, 8^ Their wide spectrum of structures and biological activities makes cyanobacterial compounds attractive as sources of drugs and drugs leads.^1, 9^

Interestingly, the molecular structures of NPs from marine cyanobacteria are distinct from those of their terrestrial and freshwater relatives;^4, 10^ they are composed of nitrogen-rich scaffolds with significant structural diversity and modifications resulting from halogenation, methylation, and oxidation.^2^ Marine cyanobacterial NPs are frequently produced by non-ribosomal peptide synthetase (NRPS), polyketide synthase (PKS), or NRPS-PKS hybrid pathways with a variety of tailoring steps.^11^ Marine cyanobacterial NPs include bioactive chemicals with diverse structures and bioactivities that could be useful in the treatment of cancer, neurological disorders, and infectious diseases;^12^ and that have anti-inflammatory properties^13^ or confer UV protection.^14, 15^

The isolation of NPs from environmental samples or cyanobacterial laboratory cultures typically yields low quantities of the compound, making full characterization of the molecules challenging. The assessment of the full potential of these molecules as drug leads often requires producing the compounds by organic synthesis.^16^ Transcriptomics and metabolomics along with genome sequence bioinformatics have also revealed that a large fraction of these BGCs are not expressed in standard laboratory cultures, leaving a substantial fraction of diverse cyanobacterial natural product compounds unexplored.^17^ Recent studies have shown that altered growth conditions can result in upregulation of some cryptic BGCs;^17, 18^ however, this approach still leaves many BGCs unexpressed in laboratory growth conditions.

Many advances have been made for the screening, detection, and identification of cyanobacterial NPs^16, 19, 20^ and their BGCs.^4, 5, 21^ However, none of these filamentous marine cyanobacteria have been genetically tractable to date. The lack of genetic methods and tools for these strains, coupled with the limited development of appropriate heterologous expression hosts, has hampered production of the larger amounts of these specialized compounds that are necessary for pharmacological studies. Investigations of the genetic underpinnings of cyanobacterial biosynthetic pathways have lagged behind those of some prominent classes of heterotrophic bacteria, such as actinobacteria and myxobacteria.^22^

Ribosomally synthesized and post-translationally modified peptides (RiPPs) including patellamide, microviridins, and mycosporine-like amino acids (MAAs) from cyanobacteria have been produced in *E. coli*.^23–26^ However, heterologous expression of NRPS, PKS, and NRPS-PKS hybrid biosynthetic pathways has proven to be more difficult.^27^ Successful production was obtained in *E. coli* upon replacement of the native promoters for lyngbyatoxin A, which is encoded by a relatively small 11.3-kb NRPS-terpene hybrid gene cluster, originally obtained from an environmental collection of *Moorena* (*Moorea*) *producens.*^28^ Two microcystin congeners, encoded by a 55-kb hybrid PKS/NRPS gene cluster from *Microcystis aeruginosa* PCC 7806, were also produced in *E. coli*.^29^ The production of lyngbyatoxin A was also attempted in *Streptomyces coelicolor* A3(2) but did not succeed,^30^ whereas the polyketide– peptide hybrid barbamide derivative 4-*O*-demethylbarbamide was produced in *Streptomyces venezuelae* but with a very low yield (<1 μg/L).^31^ Finally, the lyngbyatoxin A BGC was expressed in the cyanobacterium *Anabaena* (*Nostoc*) sp. strain PCC 7120 (hereafter *Anabaena*) and led to the production of lyngbyatoxin A with yields comparable to those of the native *M. producens*.^32^ Recently, the lyngbyatoxin pathway was further engineered in *Anabaena* to produce pendolmycin and teleocidin B-4.^33^

The development of approaches for the heterologous expression of NP pathways is important to facilitate the characterization and screening of cyanobacterial NPs for pharmaceutical applications. Strains of the genus *Moorena,* previously named *Moorea*^34^, have been found to harbor over 40 different biosynthetic gene clusters and close to 200 novel NPs have been chemically identified from *Moorena* spp.^4^ *Moorena* strains have been obtained from the photic zone of tropical marine reefs, rocks, and mangroves around the globe. There are no genetic methods for any *Moorena* strain and they grow slowly with cell division occurring only once every 6 days.^35, 36^

The *Moorena producens* strain JHB (hereafter *M. producens*), isolated from Hector’s Bay, Jamaica, carries 44 BGCs in its genome, including those encoding for production of hectochlorin, hectoramide, jamaicamide, and the recently discovered compound cryptomaldamide.^4, 16, 37^ Interestingly, cryptomaldamide is somewhat structurally similar to guadinomine B, an anti-infective compound produced by *Streptomyces* sp. K01-0509. Guadinomine B has an unusual mode of action and is a very potent inhibitor of the type III secretion system (TTSS).^38^

The putative cryptomaldamide biosynthetic pathway in *M. producens* is encoded by a 28.7-kb gene cluster (Figure 1).^16^ To facilitate studies of cryptomaldamide biosynthesis and pharmacology, we transferred the *M. producens* cryptomaldamide BGC pathway into two genetically tractable model strains of cyanobacteria, and successfully obtained cryptomaldamide production in *Anabaena*.

**Figure 1.**
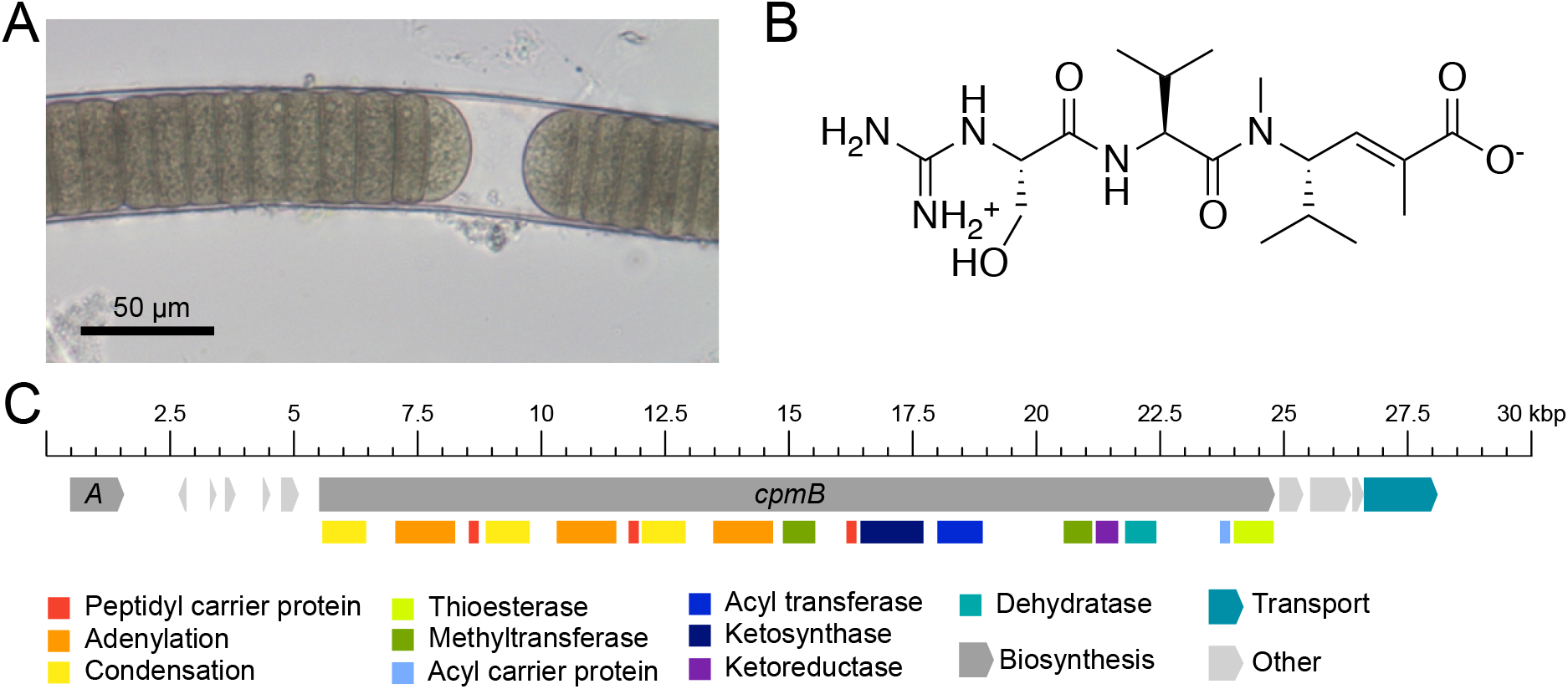
Biosynthesis of cryptomaldamide by *M. producens*. (A) Micrograph of *M. producens*. (B) Cryptomaldamide structure. (C) Putative biosynthetic gene cluster for the biosynthesis of cryptomaldamide (*cpm*) in *M. producens*. The *cpmA* gene encodes for an amidinotransferase that is believed to initiate the pathway through transfer of an amidino group from arginine to serine to produce an amidino-serine residue. The *cpmB* gene encodes a NRPS-PKS megasynthase.

## RESULTS AND DISCUSSION

### Construction of a TAR cloning plasmid and capture of the cryptomaldamide BGC

We first attempted to express the cryptomaldamide BGC in *Synechococcus elongatus* strain PCC 7942 (hereafter *S. elongatus*) because of its many advantages as a well-studied cyanobacterial model strain. *S. elongatus* is used for the study of basic biology such as its bacterial circadian clock and as a platform for synthetic biology and genetic engineering.^39, 40^ *S. elongatus* primary metabolism has been extensively studied and modeled.^41, 42^ *S. elongatus* grows rapidly and has a streamlined genome and facile genetics. It is naturally competent for DNA uptake, and large DNA fragments can be efficiently transferred by conjugation from *E. coli*.^43, 44^ Gene knockouts can be easily made and segregation can be achieved efficiently. Gene knockins and replacements can be done at native or neutral sites on the chromosome. Three neutral sites, which are sites or genes where ectopic or heterologous sequences can be inserted with no effect on known phenotypes, are commonly used for in *S. elongatus*. A large number of genetic tools are available for *S. elongatus*.^45–48^ The strain has been used as a platform for the heterologous production of various compounds including short chain alcohols, olefins, fatty acids, hydrocarbons, organic acids sugars, diols, and polyols.^49^ Recently, we reported the heterologous production of methyl branched wax esters in *S. elongatus*; this involved engineering the production of a methylmalonate precursor, the expression of the *Bacillus subtilis* promiscuous Sfp phosphopantetheinyl transferase (Sfp-PPTase), and developing a T7-polymerase expression system.^50^

To assemble the cryptomaldamide BGC from the non-genetically tractable cyanobacterium *M. producens*, we used transformation-associated recombination (TAR) in yeast (Figure 2).^51^ The cryptomaldamide BGC was amplified by PCR from genomic DNA as 3 overlapping fragments and cloned into pAM5571 (Figure 3) in *Saccharomyces cerevisiae* VL6-48N by TAR. The resulting plasmids isolated from yeast cells were screened by PCR, and positive plasmid clones were then transferred into *E. coli* and verified by restriction digests.

**Figure 2.**
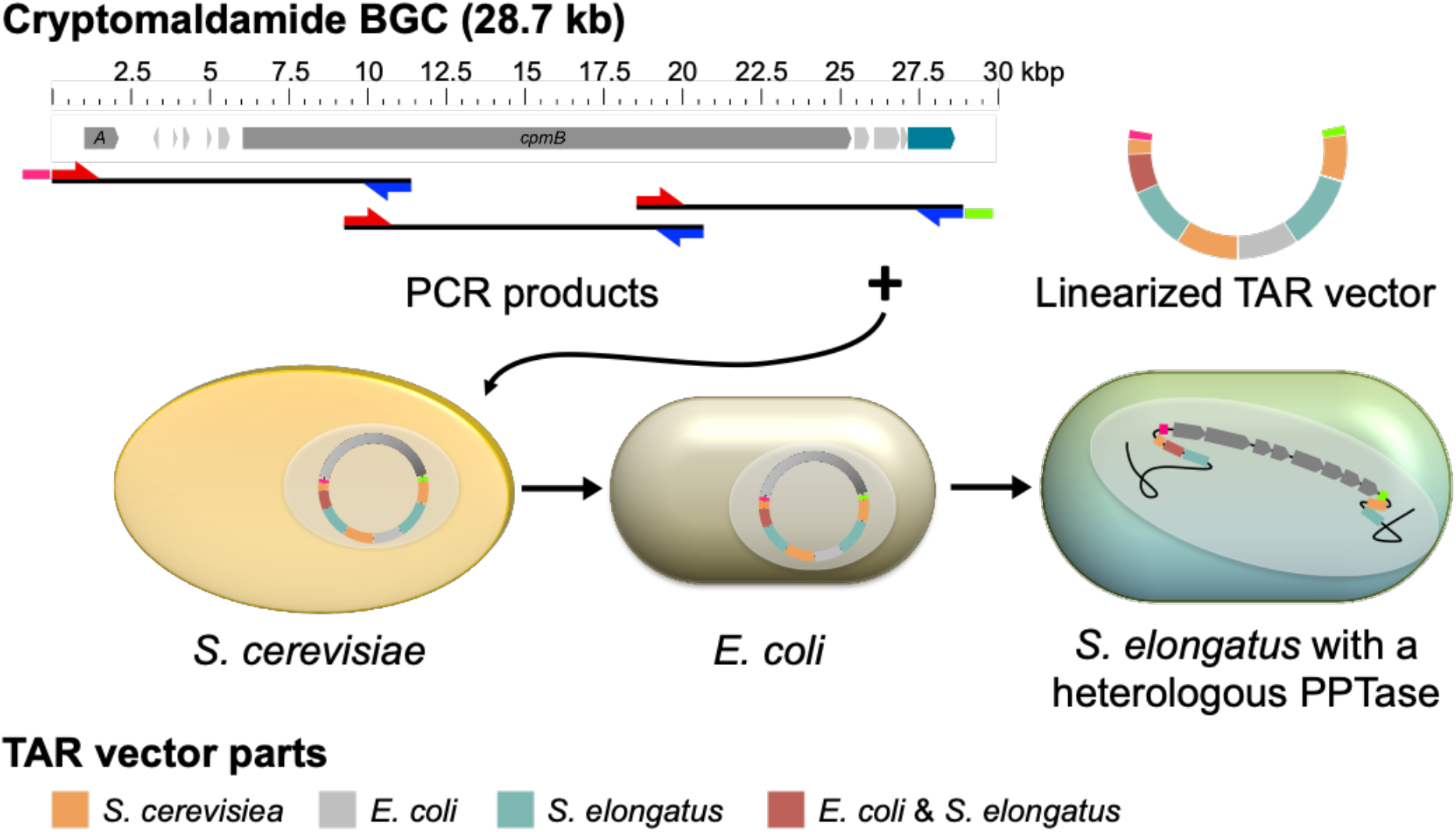
Strategy for cloning the cryptomaldamide BGC into *S. elongatus*. The cryptomaldamide BGC was amplified by PCR from genomic DNA as 3 overlapping fragments covering 28,095 bp starting 408 nucleotides upstream of the *cpmA* start codon to 47 nucleotides downstream of a multi-antimicrobial extrusion protein (MATE) efflux family protein gene. The first and last PCR products carried 40 nucleotides, pink and green dashes, that overlap with the ends, pink and green segments, of the linearized *S. elongatus* TAR cloning vector pAM5571. The 3 PCR products and pAM5571 were assembled in *S. cerevisiae* by recombination. Yeast clones containing plasmids carrying the entire BGC were identified by PCR. Positive plasmids were then transformed into *E. coli* and further verified by restriction digests with NcoI. Finally, positive plasmid clones prepared from *E. coli* were transformed into *S. elongatus*. Red arrows, forward primers; blue arrows, reverse primers.

**Figure 3.**
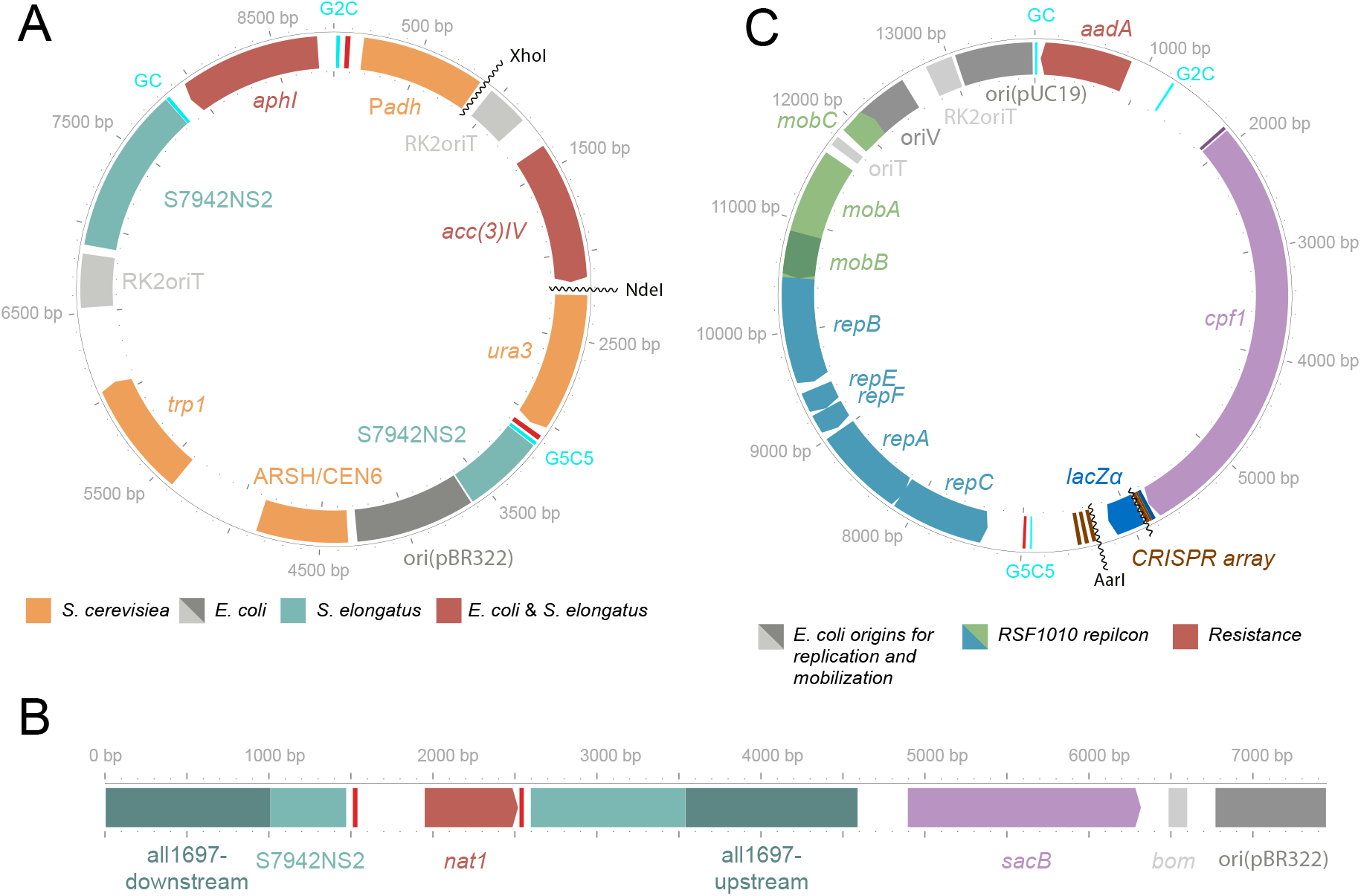
Cloning vectors and *Anabaena* chromosome engineering. (A) TAR cloning vector that comprises the following modules: (1) yeast components ARSH/CEN6 for replication and *trp1* encoding tryptophan synthetase and *ura3* encoding orotidine-5-phosphate decarboxylase (ODCase) for selection or counter selection in yeast,^54^ (2) the pBR322 origin of replication and the antibiotic resistance genes for kanamycin (*aphI*) and gentamycin (*aac(3)IV*) for replication and selection in *E. coli*, (3) the origin of transfer (RK2oriT) of the RK2 plasmid for conjugation into other microorganisms including cyanobacteria, and (4) *S. elongatus* NS2 homologous sequences for recombination into neutral site 2 (NS2). (B) Linear map of the plasmid constructed to engineer the *Anabaena* chromosome to harbor the *S. elongatus* NS2. (C) CRISPR vector that includes the *cpf1* gene and CRISPR array from pSL2680,^55^ a Sp/Sm resistance *aadA* gene, and a modified RSF1010 replicon carrying an RK2 origin of transfer and a high-copy-number *E. coli* origin of replication.^56^

### Heterologous expression of the cryptomaldamide BGC in *S. elongatus*

Initially, plasmid DNA from 4 independent positive *E. coli* clones containing the cryptomaldamide BGC were transferred by natural transformation into *S. elongatus* AMC2566. This strain has been engineered to express the promiscuous *B. subtilis* Sfp PPTase.^50^ Transcription of the BGC was evaluated and cultures were screened for the production of cryptomaldamide by LC-MS/MS. Although RT-qPCR demonstrated transcription of the BGC (Figure S1), none of the 4 strains produced cryptomaldamide at detectable levels.

Because the BGC was reconstructed using large DNA fragments produced by PCR, we were concerned that deleterious mutations may have been introduced into the BGC leading to non-functional enzymes. To overcome this potential problem, we made 64 new clones of *S. elongatus*, each carrying the BGC after transformation with a pool of 84 independent plasmid clones that had been verified by restriction digests (Figure S2). These *S. elongatus* transformants were then analyzed by MALDI analysis as previously reported for cryptomaldamide,^16^ but none produced cryptomaldamide at detectable levels. To more definitively eliminate the possibility that deleterious mutations were the cause of the failure to produce cryptomaldamide, 6 plasmid clones were sequenced. The sequencing reads were mapped onto the predicted sequence of the desired plasmid and the genetic variations were identified using the breseq program.^52^ We found that all 6 clones contained the same single nucleotide change in the *cpmA-cpmB* intergenic region (Table S1), which suggests that it may have preexisted in the gDNA for the native pathway or possibly was introduced during the early cycles of the PCR. One clone, CR92, did not carry any other mutations (Table S1). The CR92 plasmid clone, which we show below was capable of encoding for the production of cryptomaldamide in *Anabaena*, was transformed into *S. elongatus* and the resulting strain was evaluated by MALDI analysis and LC-MS/MS; however, neither cryptomaldamide nor any related compounds were produced at detectable levels.

Previous reports indicated that the cyanobacterial strain *Anabaena* PCC 7120 has an Sfp-PPTase, HetI, that is similar to the promiscuous *Bacillus subtilis* Sfp-PPTase and has high activity for carrier protein substrates of several NRPS and NRPS/PKS hybrid pathways from cyanobacteria.^32, 53^ To address the possibility that PPTase activity was the limiting factor for production of cryptomaldamide in *S. elongatus*, we expressed the *Anabaena hetI* gene from the constitutive conII promoter in the strain of *S. elongatus* carrying the cryptomaldamide pathway. However, heterologous *hetI* expression did not result in production of cryptomaldamide in *S. elongatus*, which could be caused by several problems such as inefficient translation, protein folding, or protein stability.

### Heterologous expression of the cryptomaldamide BGC in *Anabaena*

Although we did not obtain cryptomaldamide production in *S. elongatus*, we reasoned that the synthesis of functional proteins for this pathway might succeed in a different host that contained native PKS and NRPS/PKS pathways. *Anabaena* is a well-established model strain for nitrogen-fixing filamentous cyanobacteria, and it has good growth properties and well-developed genetic tools. Importantly, lyngbyatoxin A, another natural product of Hawaiian strains of *M. producens*, was successfully produced in *Anabaena* and several promoters isolated from other *M. producens* BGCs (e.g. barbamide A, curacin A, and jamaicamide) were shown to be active in *Anabaena*.^32^ In addition, the *Anabaena* genome has a low (41%) GC content that is similar to *M. producens* (44%), whereas the *S. elongatus* genome has a high GC content (55%). Finally, we reasoned that it is possible that *Anabaena* could provide a better protein-folding environment for large NP biosynthetic pathway enzymes because it has a larger genome containing several PKS and NRPS/PKS hybrid BGCs.^6^

Because the cryptomaldamide BGC was already cloned in a vector for integration into the *S. elongatus* chromosome at neutral-site 2, we realized that it would be useful for current and future experiments to place the *S. elongatus* NS2 region of homology into the *Anabaena* chromosome. Therefore, we engineered *S. elongatus* neutral-site 2 (S7942NS2) homology regions flanking an antibiotic resistance gene for nourseothricin (Nt) into the *Anabaena* chromosome at a previously identified neutral site in the all1697 gene^57^ to create *Anabaena* strain AMC2556, which carries neutral-site 2 (named A7120NS2) (Figure 3).

The plasmid from clone CR92 carrying the cryptomaldamide BGC was transferred into the AMC2556 by biparental conjugation. When a plasmid carrying homologous DNA sequences is transferred to *Anabaena*, single cross-over events that integrate the entire plasmid occur more often than double cross-over events that lose the vector sequences.^58^ In addition, like other cyanobacteria, *Anabaena* cells contain 4 to 8 copies of the chromosome and the isolation of stable mutants requires the segregation of engineered chromosomes. The selection of segregated double recombinant clones in *Anabaena* can be facilitated by a *sacB* gene carried on the plasmid backbone and selection on media containing sucrose.^58^ Here, because CR92 did not carry the *sacB* gene, we devised an alternative strategy that relied on a CRISPR/Cpf1 system shown to work well in several strains of cyanobacteria including *Anabaena*.^55, 59^ Because previously developed plasmids were not directly useable due to antibiotic incompatibilities, we constructed a CRISPR/Cpf1 module compatible with the CYANO-VECTOR platform^48^, which enabled the construction of CRISPR/Cpf1 plasmids with different antibiotic resistance genes and replicons. The CRISPR/Cpf1 module was then assembled with a spectinomycin/streptomycin resistance gene, and a modified RSF1010 replicon carrying the high copy pUC19 origin of replication,^56^ to produce pAM5572 (Figure 3). Subsequently, a guide RNA template designed to target the nourseothricin resistance (Nt^R^) gene in the A7120NS2 neutral site in strain AMC2556 was cloned into pAM5572 by golden gate cloning^60^ using 2 complementary annealed oligonucleotides to produce pAM5565. To obtain segregated double recombinant strains of *Anabaena* carrying the cryptomaldamide BGC, pAM5565 was transferred into an unsegregated strain of *Anabaena* carrying the cryptomaldamide BGC that was shown to produce cryptomaldamide in preliminary analyses. Isolated colonies were then counter-screened on nourseothricin-containing plates and sensitive clones were PCR-verified for complete segregation (Figure S3).

### Production cryptomaldamide in *Anabaena*

#### Detection of cryptomaldamide in the cell biomass and the growth medium

Before we obtained segregated clones, three independent colonies (AMC2560, AMC2561, AMC2562) of *Anabaena* carrying the cryptomaldamide BGC were selected, analyzed by PCR for the presence of the BGC, and then grown in cultures to determine if they produced cryptomaldamide. These clones and AMC2556 as a negative control were grown for 10 days in 150 mL BG-11 medium bubbled with air to an OD_750_ of approximately 0.8. Extracts of the cell biomass and the growth medium were analyzed by LC-MS/MS and both showed a peak matching the retention time and molecular ion *m/z* value of cryptomaldamide ([M + H]^+^ *m/z* 400) obtained from *M. producens* (Figure 4A and B). Moreover, the MS/MS fragmentation pattern matched the previously reported fragmentation pattern of cryptomaldamide (Figure 4B).^16^ These data provided preliminary confirmation of heterologous production of cryptomaldamide in *Anabaena*.

**Figure 4.**
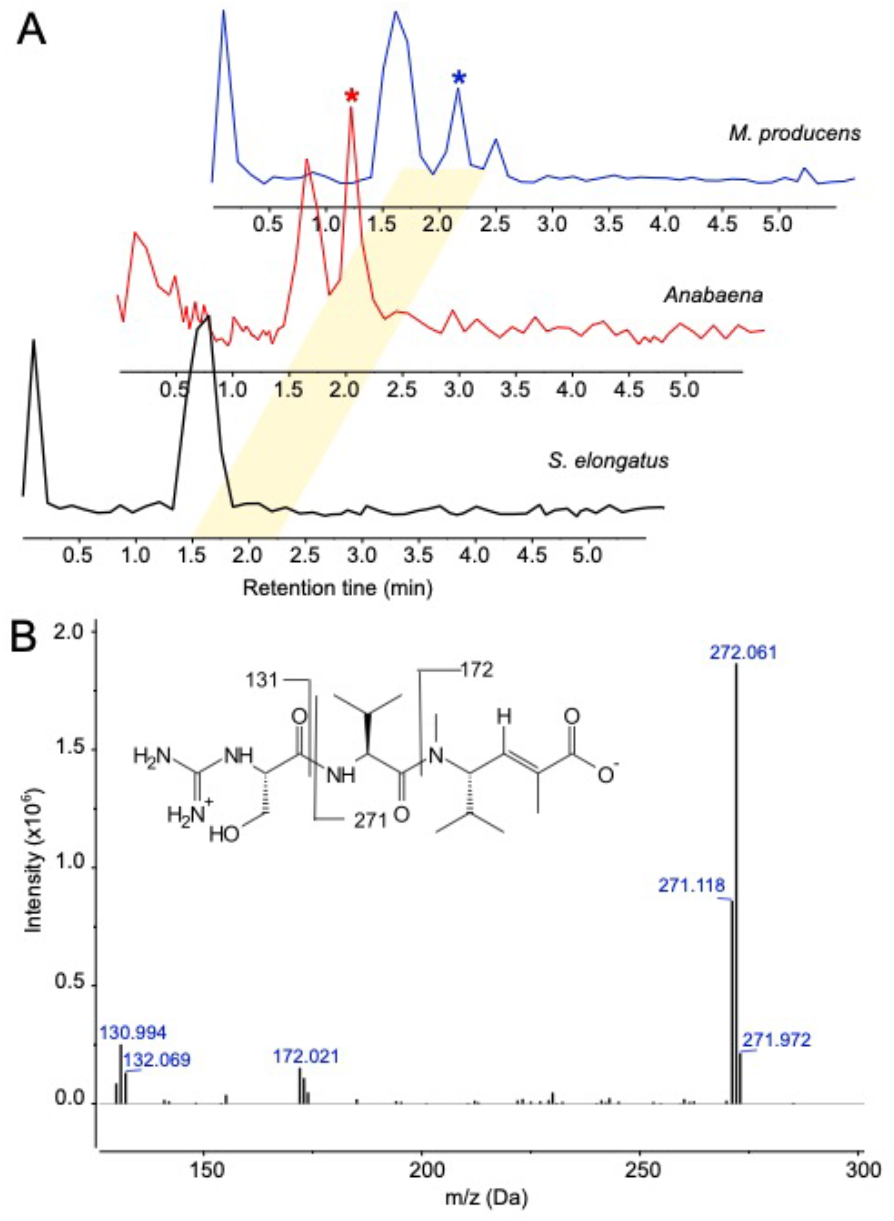
Detection of cryptomaldamide. (A) Representative LC-MS Total Ion Chromatograms (TICs) of the extracts obtained from the native *M. producens* strain and heterologous hosts *Anabaena* and *S. elongatus* carrying the cryptomaldamide BGC. Cryptomaldamide peaks are indicated with asterisks (*). (B) Representative tandem mass spectrum indicating *m/z* fragmentations of cryptomaldamide isolated from a culture of *Anabaena* that expresses the cryptomaldamide BGC.

#### Isolation of cryptomaldamide and structure confirmation

Two segregated double recombinant clones of *Anabaena* (AMC2564 and AMC2565) were each grown as 1.5-L cultures in a 3% CO_2_ atmosphere for about 3 weeks until they reached an OD_750_ of 2.3-2.6. To simplify downstream purification steps, the cell biomass was removed from the growth medium by centrifugation. Half of each culture supernatant was concentrated by evaporation and the residue was dissolved in MeOH and then dried, resulting in 28 mg of crude organic extract. This extract was fractionated by RP-HPLC to produce 14.6 mg of a pure, white amorphous solid. The 500 MHz ^1^H NMR spectrum of this purified compound in DMSO-*d*_6_ matched the reported proton assignments for cryptomaldamide isolated from *M. producens* (Table S2).^16^ Additionally, two singlets at δ_H_ 8.03 - 8.09 and a singlet at δ_H_ 7.78 were indicative of the presence of a monosubstituted guanidine group. Taken together, these data confirmed that cryptomaldamide was being produced in the heterologous host *Anabaena*.

#### Production of cryptomaldamide over time and under different culture conditions

One of the prominent motivations to produce NPs in a heterologous host is to increase the supply of compounds for pharmacological testing. Therefore, we evaluated culture conditions, growth phase, and substrate availability to determine the effects on the amount of cryptomaldamide produced by *Anabaena*. To quantify cryptomaldamide production, we created a standard concentration curve using purified cryptomaldamide that had a limit of detection of 1.5 ng and was linear up to 1.56 μg (Figure S4).

To determine the production of cryptomaldamide over time, *Anabaena* AMC2564 and AMC2565 were grown in 2-L cultures in a 3% CO_2_ atmosphere starting at an OD of 0.05, and samples were collected after one day (OD_750_ ~0.2), 1 week (OD_750_ ~0.6), and 3 weeks (OD_750_ ~1.5). Cryptomaldamide was measured from the culture biomass and from the medium (Figure 5A). Cryptomaldamide accumulated in these larger slow-growing cultures over time and reached a total concentration of 25.9 ± 3.6 mg/L after 3 weeks. The concentration of cryptomaldamide in the biomass increased over time, starting at less than 1.5 ± 1.0 mg/g dry weight and increasing to 15.3 ± 2.4 mg/g biomass dry weight after 3 weeks (Figure 5B). In contrast, the amount of cryptomaldamide in the medium was directly correlated with the cell biomass at each time point and remained at a little over 5 mg/g biomass dry weight. Cryptomaldamide accumulated in *Anabaena* cells to higher levels than was released into the medium. Additional investigation will be required to understand whether the cryptomaldamide in the medium is actively exported (for example using the MATE protein encoded in the BGC), passively diffuses into the medium, or is released by lysis of some of the cells. Interestingly, the amounts of cryptomaldamide in *Anabaena* biomass were approximately 20-fold higher compared to *M. producens*, which contains 0.7 mg/g dry weight.^16^ It took about 3 weeks to harvest 1.2 gram dry weight of cell biomass per liter from each of the engineered *Anabaena* AMC2564 and AMC2565 cultures; based on previous studies, we estimate that it would take about 9 weeks, 3 times longer, to harvest the same amount of biomass of *M. producens* from laboratory cultures.^61^

**Figure 5.**
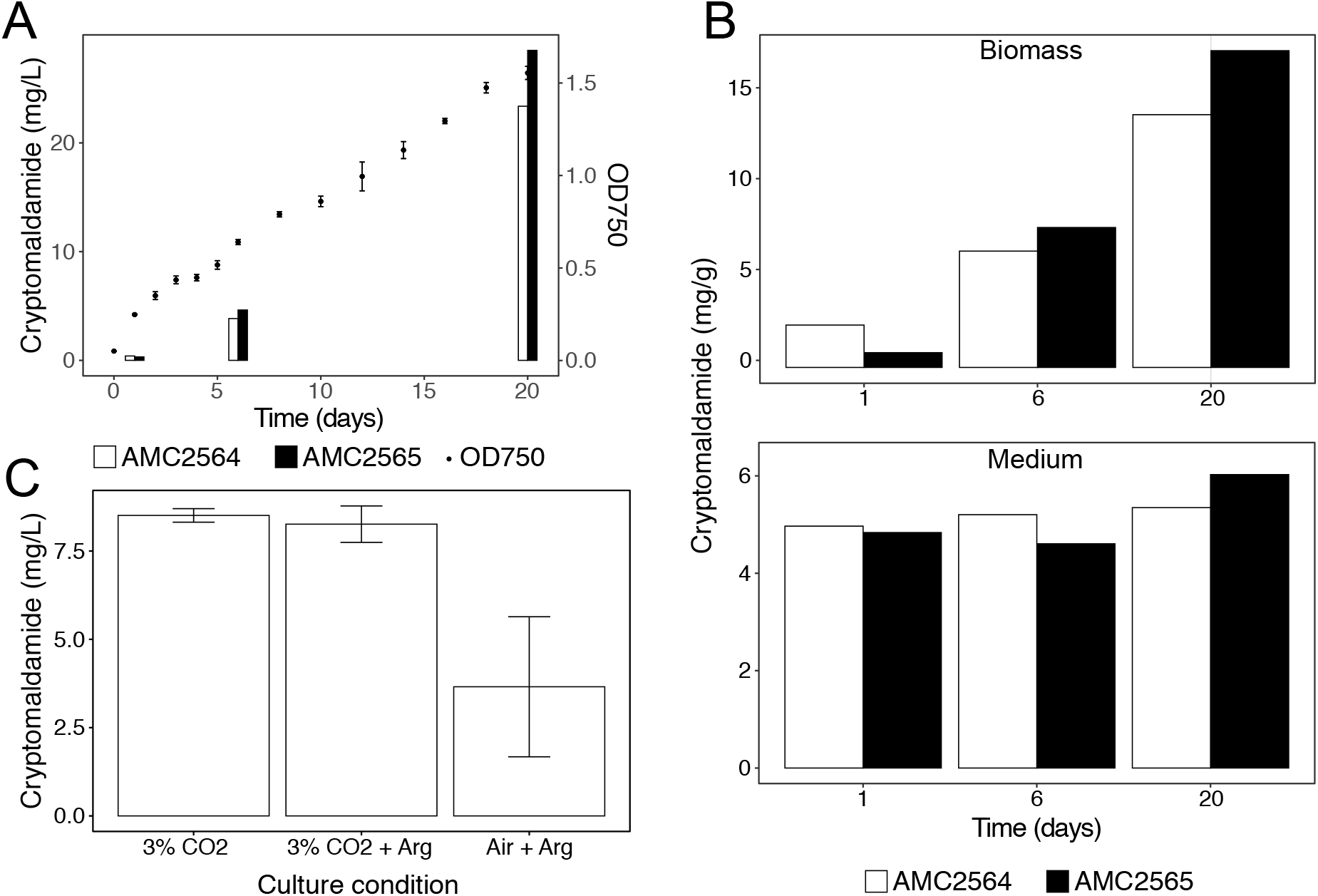
Production of cryptomaldamide in *Anabaena*. (A) Total cryptomaldamide concentrations (bars) at 3 time points (1, 6, and 20 days) obtained for cultures (cell biomass + growth medium) of two independent *Anabaena* clones (AMC2564 and AMC2565) that contain the cryptomaldamide BGC, and cell density (dots, OD_750_) over a 20-day time course. 2-L cultures were grown in 2.8-L Fernbach flasks in a 3% CO_2_ atmosphere. The OD_750_ values are the mean values ± standard deviations for the 2 cultures. OD_750_ was measured every day for the first 6 days and every 2 days for the rest of the time course. (B) Amount of cryptomaldamide in the biomass and the medium normalized to cell biomass dry weight. The cells and medium were collected on days 1, 6, and 20. (C) Concentration of cryptomaldamide in 50-mL cultures of AMC2565 grown in 125-mL flasks in a 3% CO_2_ atmosphere with and without 5 mM arginine and grown in air without supplemental CO_2_. Cultures were grown until the cells reached an OD_750_ of approximately 1.5. Cryptomaldamide concentrations were normalized to cell density by dividing by the culture OD_750_. The experiment was carried out in triplicate for each condition, and cryptomaldamide concentrations are shown as mean values ± standard deviations.

The first residue to be incorporated by the NRPS portion of the cryptomaldamide biosynthetic pathway is an amidino-serine residue. The amidinotransferase, CpmA, is proposed to transfer an amidino group from arginine to serine to form this amidino-serine residue.^16^ Therefore, we investigated if the addition of 5 mM arginine to the medium could increase cryptomaldamide production; however, the production of cryptomaldamide was unchanged when the engineered *Anabaena* strain AMC2565 was grown in a 3% CO_2_ atmosphere with 5 mM arginine (Figure 5C). Therefore, arginine is likely not a limiting substrate for the production of cryptomaldamide in *Anabaena*; this may be because cyanobacteria store nitrogen as cyanophycin, a co-polymer of aspartate and arginine,^62^ and cells may regulate arginine levels to maintain its availability. For cultures grown in a 3% CO_2_ atmosphere, the 2-L slow-growing cultures (Figure 5A) produced more cryptomaldamide per volume of culture than did the faster-growing 50-mL cultures (Figure 5C, left bar) at similar cell densities. This indicates that in the large cultures, the cells continue to produce and accumulate cryptomaldamide after they have become light-limited for rapid growth.

Cyanobacteria fix CO_2_ via photosynthesis for growth, and several studies have shown that higher growth rates, increased maximum cells densities, and upregulation of secondary metabolite pathways occur with an increased partial pressure of CO_2_.^18, 63–65^ Therefore, we compared the amount of cryptomaldamide produced in cultures grown in air with cultures in grown in a chamber with a 3% CO_2_ atmosphere. The cultures grown in air took 4 weeks to reach an OD_750_ of ~1.5, whereas the cultures grown in 3% CO_2_ grew faster and took only one week to reach the same density. Both cultures were supplemented with arginine, but as stated above, we found that arginine addition does not affect cryptomaldamide production. The production levels of cryptomaldamide in the slower-growing cultures in air were less than half that of the faster-growing cultures in 3% CO_2_ (Figure 5C) showing that incubating cultures with supplemental CO_2_ stimulates cryptomaldamide production.

### Bioactivity testing of cryptomaldamide produced in *Anabaena*

Heterologous production of cryptomaldamide in *Anabaena* increased its availability for biological activity testing. Inspired by recent discoveries of novel NPs with guanidine groups that have antimicrobial activity,^66^ growth inhibition assays with cryptomaldamide against *Candida albicans*, *Bacillus subtilis*, and *Pseudomonas aeruginosa* were performed. However, there was no growth inhibition by cryptomaldamide at amounts up to 0.5 mg per disc (data not shown). Similarly, cryptomaldamide was previously reported to have no effect against H460 human lung cancer cells and had no blocking effects of the mammalian voltage gated sodium channel Nav1.4.^16^

## Conclusions

The marine environment provides a particularly rich source of biologically active NPs as well as their associated BGCs.^67^ Hundreds of PKS/NRPS BGCs have been identified in cyanobacterial genomes^21^, including marine strains such as *M. producens*.^4^ However, in laboratory cultures, the encoded NPs are often produced in small quantities, and moreover, many of the BGCs are silent.^68^ Genetic studies of these BGCs would enable further interrogation of their regulation and the biosynthetic mechanisms that underlie the production of specific compounds; however, most cyanobacterial strains, including marine “super producer” strains, are not genetically tractable.^56^ As is true for other microorganisms, the heterologous expression of cyanobacterial NPs is therefore a promising approach to answer such limitations.^22^

The polyketide–peptide hybrid natural product cryptomaldamide was discovered in the marine cyanobacterium *M. producens* using MALDI analysis and characterized by 2D NMR and other spectroscopic and chromatographic methods.^16^ By heterologous expression in *Anabaena*, we have experimentally confirmed that a putative 28.7-kb BGC proposed by Kinnel et al.^16^ to encode for cryptomaldamide biosynthesis is responsible for its production. These results further validate *Anabaena* as a heterologous platform for the production of cyanobacterial NPs, as was previously demonstrated for the production of the non-ribosomal peptide-terpene compound lyngbyatoxin A in *Anabaena*.^32^

Many drugs and drug leads are NPs, NP derivatives, or NP-inspired molecules.^9^ Actinobacteria and more recently myxobacteria from terrestrial and aquatic ecosystems have been important sources of NPs with a broad range of biological activities.^69, 70^ Marine cyanobacteria are also a substantial source of NPs. Hundreds of NP compounds have been identified from cyanobacteria but their exploitation as drugs is still largely untapped because of limited availability of the compounds. The mechanisms of action and applications of only a few cyanobacterial NPs have been more deeply investigated. These include a few potent cancer cell cytotoxins such as curacin A, which is a microtubule polymerization inhibitor, apratoxin A, which prevents the biogenesis of secretory and membrane proteins, and dolastatin 10 that has been modified to be the warhead of an antibody drug conjugate that is FDA approved.^71, 72^ ^73, 74^ Indeed, the slow growth rate and lack of genetic tractability of marine cyanobacteria have limited the development of their NPs by the pharmaceutical industry. The heterologous expression of NP BGCs from marine cyanobacteria in *Anabaena* is an important method for the discovery and development of valuable NPs. The well-developed genetic methods available for *Anabaena* will facilitate a better understanding of NP gene regulation and evolution, and enable studies of enzymatic mechanisms and biosynthesis. The success expressing cryptomaldamide in a heterologous host is an important step in developing high-throughput technologies for cyanobacterial NP exploration, production, and downstream analysis.

## METHODS

### Plasmid constructions

Plasmids and oligonucleotides used in this study are listed in Table S3 and S4, respectively.

PCR amplifications were carried out with Q5 High-Fidelity DNA polymerase (New England BioLabs) according to the manufacturer’s instructions. Plasmid preparations were performed using the QIAprep Spin Miniprep Kit (Qiagen). Restriction digests followed the supplier’s recommendations but with longer incubation times to assure complete digests. DNA purification/concentration following PCR and restriction digests were performed with DNA Clean & Concentrator TM-5 (Zymo). Nucleic acid concentrations were measured with a NanoDrop 2000c spectrophotometer. Cloning in *E. coli* was carried out by restriction/ligation with NEB Quick Ligase following the manufacturer instructions or by Gibson assembly using the GeneArt Seamless Cloning and Assembly Kit (Thermo Fisher) as described previously.^48^ Transformation-associated recombination cloning in the yeast *Saccharomyces cerevisiae* was performed using 500 ng of plasmid backbone linearized with XhoI and NdeI and PCR products in equimolar ratios following previously published protocols.^75, 76^

To make pAM5273, pCVD022 was digested with PciI and AflII, the yeast element ARSH/CEN6 was PCR amplified from pCAP03-acc(3)-IV with the primer pair S7942NS-Yeast-F/S7942NS-Yeast-R, then the resulting DNA fragments were assembled by seamless cloning. To make pAM5276, pCVD015 was digested with SwaI, a DNA fragment that contains the *adh* promoter and the *ura3* gene was PCR amplified from pCAP03-*acc(3)-IV*, then the resulting DNA fragments were assembled by seamless cloning. To make pAM5277, the EcoRV site in the *ura3* gene was edited by quick change PCR of pAM5276 using the complementary primers ura3-t189c-F and ura3-t189c-R. To make pAM5601, pAM5372 was digested with EcoRI and SbfI, the *hetI* (all5359) gene was PCR amplified from *Anabaena* PCC 7120 gDNA with the primer pair pAM5372_A7120_hetI_2F/pAM5372_A7120_hetI_583R, then the resulting DNA fragments were assembled by seamless cloning. To make pAM5564, pAM5571 was digested with XhoI and NdeI, the cryptomaldamide BGC was PCR amplified from *M. producens* gDNA as 3 fragments with the primer pairs P100F/P100R, P101F/P101R, P102F/P102R, then the resulting DNA fragments were assembled by TAR cloning in *S. cerevisiae*. To make pAM5565, pAM5572 was digested with AarI, the DNA fragment coding for the crRNA was obtained by annealing the phosphorylated oligonucleotides NT_U248F and NT_U228R, then cloned into pAM5572 by ligation. To make pAM5569, pER015 was digested with XbaI and NheI, a DNA fragment that contains the *S. elongatus* NS2 homology sequences flanking a nourseothricin resistance gene was PCR amplified from pAM5544 with the primer pairs A7120NS2xS7942NS2LAF/A7120NS2xS7942NS2RAR, and then the resulting DNA fragments were assembled by seamless cloning. To make pAM5571, pAM5273 was digested with ZraI to obtain DNA a fragment that contains the *S. elongatus* NS2 homology sequences and yeast elements, pCVD003 was digested with EcoRV to obtain the *aphI* gene, and pAM5277 was digested with EcoRV to obtain the *Padh-ura3* cloning module, and then the resulting DNA fragments were assembled by seamless cloning. To make pAM5572, pAM5406 was digested with ZraI to obtain the modified RSF1010 replicon, pCVD002 was digested with EcoRV to obtain the *aadA* gene, and pAM5600 was digested with EcoRV to obtain the *cpf1*/CRISPR module, and then the resulting DNA fragments were assembled by seamless cloning. To make pAM5600, the *cpf1*/CRISPR system from pSL2680 was digested with PstI and SalI, the backbone of pCVD015 was PCR amplified from pCVD015 with the primer pair pCVD015_1684F/pCVD015_3464R and digested with PstI and SalI, and then the resulting DNA fragments were assembled by seamless cloning.

Plasmid constructs were verified by restriction digests, and DNA fragments produced by PCR were sequence-verified by Sanger sequencing. Six independent plasmid clones carrying the cryptomaldamide BGC were sequenced entirely by next-generation sequencing on a MiSeq platform.

### Strain construction and culture conditions

Cyanobacterial strains used in this study are listed in Table S5.

Plasmid DNA was introduced into *S. elongatus* AMC2302, which has been cured of the endogenous pANS plasmid, by natural transformation using standard protocols.^46^ Recombinant DNA carrying the cryptomaldamide BGC and an antibiotic resistance marker were integrated into the chromosome at neutral site 2 (NS2).^46^ Recombinant DNA carrying an Sfp-PPTase and an antibiotic resistance gene were integrated into the chromosome at NS3^77^ or carried on a replicative plasmid derived from the *S. elongatus* small plasmid pANS.^48^ For *Anabaena*, recombinant DNA was introduced into cells by biparental conjugations from *E. coli* following published protocols.^78, 79^

To construct *Anabaena* strain AMC2556, which contains the *S. elongatus* NS2 neutral site in the chromosome, recombinant DNA carrying *S. elongatus* NS2 homology regions flanking a Nt^R^ gene (pAM5569) was integrated into the all1697 gene^57^ in the *Anabaena* PCC 7120 chromosome. We named this neutral site A7120NS2. To obtain segregated double recombinant strains after conjugation of pAM5569, which carries a *sacB* gene on its backbone, isolated colonies were pooled, and several dilutions of the mixture were plated onto BG-11 plates supplemented with 5% sucrose. Several isolated colonies were further grown as small patches on fresh plates and segregation was verified by PCR. One segregated strain was archived as strain AMC2556.

Plasmid pAM5564, which carries the cryptomaldamide BGC, was conjugated into *Anabaena* AMC2556 and 3 independent neomycin resistant (Nm^R^) clones were selected: AMC2560, AMC2561, and AMC2562. To obtain segregated double recombinant clones of *Anabaena* carrying the cryptomaldamide BGC in the A7120NS2 neutral site, pAM5565, which was designed to cleave the Nt^R^ gene in AMC2556 and providing spectinomycin and streptomycin resistance (Sp^R^ + Sm^R^), was conjugated into AMC2560. Then, isolated colonies (Nm^R^, Sp^R^ + Sm^R^) were counter screened on plates with Nt, and 3 clones that did not grow in the presence of Nt were grown as small cultures in BG-11 with Nm. Finally, to obtain strains that had lost pAM5565, aliquots of these cultures were spread on plates with Nm, isolated colonies were counter screened on plates with Sp + Sm and for each clone, one colony that did not grow in the presence of Sp + Sm was verified by PCR (Figure S3).

*E. coli* strains were grown at 37 °C in LB broth or on agar plates supplemented with appropriate antibiotics. *S. cerevisiae* VL6-48N strains were grown at 30 °C in YPD medium supplemented with 100 mg/L adenine in broth culture or on agar (2%) plates. Cyanobacterial strains were grown in BG-11 medium^80^ as liquid cultures at 30 °C with continuous shaking or on agar plates (40 mL, 1.5% agar). *S. elongatus* was grown with continuous illumination of 300 μmol photons m^−2^ s^−1^ and *Anabaena* was grown with 70 μmol photons m^−2^ s^−1^. Cultures were grown in air with fluorescent cool white bulbs as the light source. Culture media for recombinant cyanobacterial strains were supplemented with appropriate antibiotics: for *S. elongatus*, kanamycin (5 μg/mL) and chloramphenicol (7.5 μg/mL); and for *Anabaena*, neomycin (12.5-25 μg/mL) and nourseothricin (25 μg/mL).

### RT-qPCR

Four independent cultures of *S. elongatus* carrying the cryptomaldamide pathway and the host strain AMC2556 where grown until they reached an OD_750_ of ~0.5. Cell collection, RNA extraction, cDNA synthesis, and qPCR were performed as described previously.^43^ Primer sequences and target genes are listed in Table S4.

### MALDI analysis of*S. elongatus* cultures

Cell pellets collected from 1 mL of liquid culture or scraped from a small patch (~0.5 cm^2^) of cells grown on agar plates were resuspended in 50 μL of BG-11 medium. A 0.5 μL aliquot of the suspended cells were then mixed with 0.5 μL of matrix solution (acetonitrile(ACN): trifluoroacetic acid(TFA) 78:0.1 saturated with universal matrix from Sigma) onto a MALDI MSP 96 anchor plate (Bruker Daltonics). The plate was then air-dried for 30 minutes at room temperature and then analyzed by MALDI-TOF mass spectrometry on a Bruker Daltonics Microflex system. The data were analyzed with the MALDIquant and MALDIquantForeign packages using custom scripts written in R.^81^

### Extraction, identification, characterization, and quantification of cryptomaldamide

#### Preliminary identification

Cultures of *S. elongatus* and *Anabaena* were grown in 150 mL of medium in 250-mL Falcon tissue culture flasks bubbled with air (0.1 L/min). Cultures were typically started at an OD_750_ of 0.05-0.1 and grown until they reached an OD_750_ of ~1. To collect samples, cells were pelleted by centrifugation at 4,500 g and the supernatants and cell pellets were frozen at −80°C or processed immediately for chemical analysis. The cell pellets were extracted in 100% ethanol 3 times, then the crude extract was evaporated and redissolved in methanol for LC-MS analysis. The growth medium was liquid-liquid extracted three times with 250 ml of n-butanol and both the aqueous layer and organic layers were evaporated and dissolved into methanol for LC-MS analyses.

#### Isolation and characterization

To obtain larger amounts of cryptomaldamide from *Anabaena* AMC2464 and AMC2465, these strains were grown as 1.5-L cultures in 2.8-L Fernbach flasks on an orbital shaker at 125 rpm in a chamber with a 3% CO_2_ atmosphere. The cultures were inoculated at an OD_750_ of 0.02 and grown for 23 days until they reached an OD_750_ of 2.3 and 2.6. As the cultures became denser, the light intensity was progressively increased from 40 to 200 μmol photon m^−2^ s^−1^. The light source was natural white light LED strips (4000 – 4500 K).

Both cultures were centrifuged, and the growth medium was evaporated to give a crude product composed of salts and organic compounds. This crude product was dissolved in methanol and centrifuged for 5 minutes at 12,000 rpm. The supernatant was then subjected to purification using semi-preparative HPLC with a reverse phase gradient (5% ACN/H_2_O to 14.1% ACN/H_2_O for 6 minutes followed by an isocratic phase at 14.1% ACN/H_2_O for 6.75 minutes, at which time cryptomaldamide eluted as a single peak (Phenomenex Aqua Kinetex 5μ C18 125Å, 250 × 4.00 mm, 2.5 mL/min). Pure cryptomaldamide was analyzed by ^1^H NMR using a JEOL ECZ500R NMR operating at 500 MHz with the samples dissolved in DMSO-*d*_6_.

#### Quantification of cryptomaldamide

To determine the amount of cryptomaldamide produced over time, AMC2464 and AMC2465 were grown as described above except both strains were grown as 2 L cultures with a light intensity of 80 μmol photon m^−2^ s^−1^. At three time points, 200 mL aliquots were collected, the cells were collected by centrifugation and placed at −80 °C. The growth medium supernatant was kept at 4 °C in the dark until analysis. A 1 mL aliquot of the growth medium was partially cleaned by purification over C-18 SPE columns using 1 additional milliliter of MeOH to elute samples directly into LC-MS vials. LC-MS analyses were then conducted in triplicate for each vial. The cell pellets were lyophilized and weighed, and then extracted 3 times using 100% ethanol. The solvent in these crude extracts was evaporated, and the samples redissolved in methanol for LC-MS analyses using a Finnigan LCQ Linear Ion Trap LC/MS/MS instrument.

To determine the amount of cryptomaldamide produced under the different culture conditions, AMC2465 was grown as 50 mL cultures with or without 5 mM arginine in 125 mL flasks. One set of cultures was grown in air and another set of cultures was grown in a 3% CO_2_ atmosphere. All cultures were grown in triplicate. Preliminary experiments indicated that one freeze/thaw cycle of a culture of *Anabaena* led to the complete release of cryptomaldamide into the medium from the lysed cells, leaving no trace of cryptomaldamide in the biomass. Therefore, all culture samples were frozen and at the time of analysis, the culture samples were thawed, and the cell debris was removed by centrifugation. The supernatant was then processed similarly to the growth medium as described above for the LC-MS analyses.

Cryptomaldamide was quantified in the samples using a standard curve. The protonated molecular ion peak [M + H]^+^ at *m/z* 400.25 was used for quantification. A series of 1:2 dilutions starting with 50 μg of cryptomaldamide down to a 1.5 ng were analyzed by LC-MS and the area under the curve (AUC) for each sample was measured. Serial dilutions and LC-MS analyses were performed in duplicate and a linear regression was used to fit a standard curve to the averaged values within the limits of linearity (Figure S4).

#### General procedures

Chemical reagents were purchased from Acros, Fluka, Sigma-Aldrich, or TCI. Deuterated NMR solvents were purchased from Cambridge Isotope Laboratories. ^1^H NMR spectra were collected on a JEOL ECZ 500 NMR spectrometer equipped with a 3 mm inverse detection probe. NMR spectra were referenced to residual solvent DMSO signals (δ H 2.50 ppm and δ C 39.52 ppm as internal standards). The NMR spectra were processed using MestReNova (Mnova 12.0, Mestrelab Research). Each crude and pure sample was injected and analyzed via LC-MS/MS on a Thermo Finnigan Surveyor Autosampler-Plus/LC-MS/MS/PDA-Plus system coupled to a Thermo Finnigan LCQ Advantage Max mass spectrometer with a 10 minute gradient of 30 − 100% CH_3_CN in water with 0.1% formic acid in positive mode (Kinetex 5μ C18 100Å, 100 × 4.60 mm, 0.6 mL/min). The ion trap mass spectrometry raw data (.RAW) were converted to the *m/z* extensible markup language (.mzXML) with MSConvert (v 3.0.19) and uploaded to GNPS.^82^ Spectral library search was performed against available public libraries and NIST17. The spectra for cryptomaldamide were annotated in the GNPS spectral library (https://gnps.ucsd.edu/) (accession number: CCMSLIB00005724004).

### Biological assays

*Bacillus subtilis*, *Pseudomonas aeruginosa*, and *Candida albicans* were streaked on LB agar or Sabouraud dextrose (SD) agar (for *C. albicans*). After 24 hours, 3 isolated colonies from each plate were grown in LB or SD broth until they reached 0.5 McFarland turbidity standard. Microbial suspensions (25 μL per plate) were then spread on LB or SD agar plates. Six 6 mm Whatman paper disks containing purified cryptomaldamide or solvent controls were evenly distributed onto each plate. The paper disks were loaded with 50, 200, or 500 μg of cryptomaldamide from a 0.5 mg/mL stock solution; a maximum of 200 μL were added at a time and dried before a subsequent 200 μL volume or smaller was added. All the paper disks were dried in a fume hood before application to the test plates. Disks treated with pure MeOH or no treatment at all were used as negative controls. BD BBL Sensi Discs with 10 μg of streptomycin were used as positive controls. The plates were incubated at 30°C for 24 hours at which time zones of inhibition were measured as the diameter of the ring around the disk where microbial growth was absent.

## AUTHOR INFORMATION

### Author Contribution

A.T., J.W.G., L.G., and W.H.G. conceived the project. A.T., B.D., B.A., and N.A.M. constructed the recombinant plasmids and strains. A.T. and N.A.M. performed MALDI analyses. A.E. and R.R. performed LC-MS/MS and NMR analyses. A.T. performed RT-qPCR experiments. A.E. performed the bioassays. T.F.L. and N.A.M. provided the BGC identification. P.C.D. provided laboratory support for MALDI analysis. R.S. provided MALDI analytical scripts. A.T., A.E., J.W.G, L.G., and W.H.G. analyzed the data and wrote the paper, which was reviewed and edited by all authors.

## ACKNOWLEDGMENTS

We thank K. Jepsen at the UC San Diego IGM sequencing facility for technical support. We thank B.S. Moore for providing pCAP03 and *S. cerevisiae* VL6-48N, and X. Tang for protocols and tips on TAR cloning. We thank E. Glukhov for maintaining the laboratory culture of *M. producens*. We thank A. M. Caraballo for assistance with MALDI analyses. We thank T. Gilderman, D. Genuário, and C. Peterson for assistance with plasmid and strain construction, and S.S. Golden for general laboratory financial support. Funding was provided by the National Institute of General Medical Sciences of the National Institutes of Health under award number R01GM118815 (J.W.G., L.G., and W.H.G.) and the Department of Energy under award number DE-EE0007094 (R.S.). The content is solely the responsibility of the authors and does not necessarily represent the official views of the National Institutes of Health.

