## Supporting Information for "Heterologous expression of cryptomaldamide in a cyanobacterial host"

### FIGURES

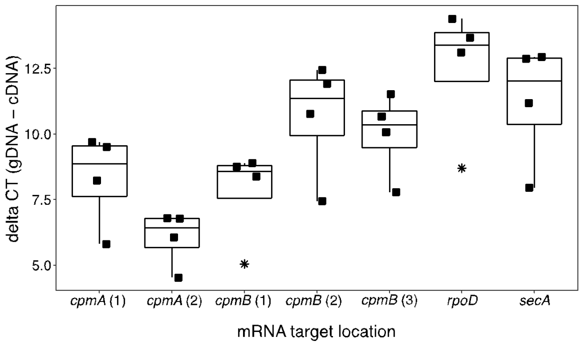


**Figure S1. Transcription of the cryptomaldamide BGC in *S. elongatus.*** Boxplot of RT-qPCR delta CT values of the *cpmA* and *cpmB* genes in the cryptomaldamide pathway and control *S. elongatus rpoD* and *secA* housekeeping genes. Delta CT values are the CT values for contaminant gDNA minus the CT values for cDNA. The CT values were normalized with the CT values obtained for the 16S rRNA in each sample. The box indicates the inter-quartile range (IQR); the center line indicates the median; the whiskers extend from the box hinges to the minimal and maximal values at most 1.5 * IQR; n = 4, outlier data points are marked as an asterisk.

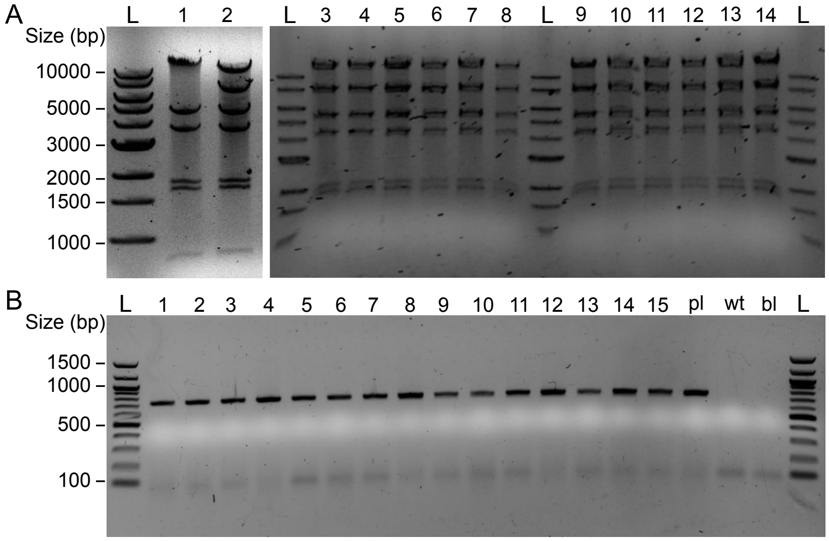


**Figure S2. Efficient capture of the cryptomaldamide BGC from PCR products in pAM5571 using TAR cloning in *S. cerevisiae* and transformation at NS2 in the *S. elongatus* chromosome.** A. Examples of NcoI restriction digests of plasmid DNA extracted from individual *E. coli* clones after TAR cloning in *S. cerevisiae* and retransformation into *E. coli*. A large majority of the clones screened by restriction digest, 84 out of 96, displayed the expected restriction pattern (expected band sizes: 13446, 7902, 5440, 4195, 1970, 1804, 805 bp). B. PCRs targeting the cryptomaldamide BGC carried in individual colonies of *S. elongatus* after transformation with pooled plasmid DNA from the 84 positive cryptomaldamide BGC clones. Every colony that was screened carried the cryptomaldamide BGC (expected PCR product band size: 773 bp) and 80% of the clones were double recombinants that had lost the suicide vector backbone (data not shown). Individual clones are indicated with a number; pl, plasmid; wt, wild type; bl, blank; L, 1-kb size-marker ladder.

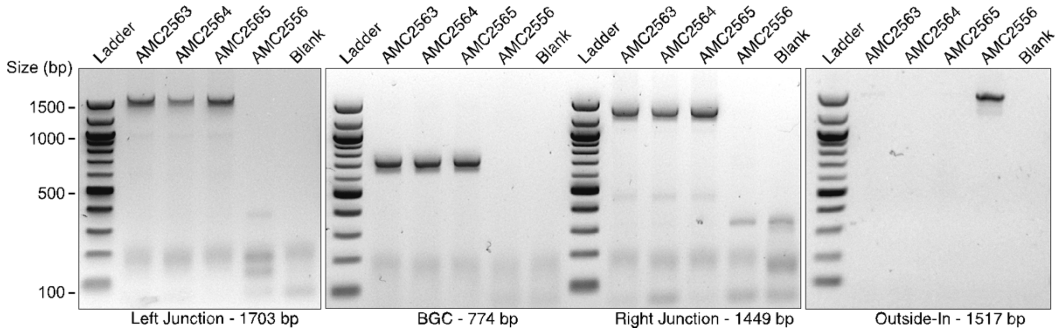


**Figure S3. PCR assay for the cryptomaldamide BGC in recombinant strains of *Anabaena* and for segregation in double recombinants.** The strains AMC2563, AMC2564, and AMC2565 carrying the cryptomaldamide BGC were compared to the NS2 platform strain AMC2556. The leftmost 3 sets of PCRs targeted the *cpmB* gene in the BGC including the left and right junctions at the chromosomal insertion sites. The last set of PCRs used primers on both sides of the insertion site to determine if the strains were fully segregated. A faint band was detected in AMC2563, which indicates that this strain was not completely segregated. The expected PCR product band sizes are listed at the bottom of each set of PCRs.

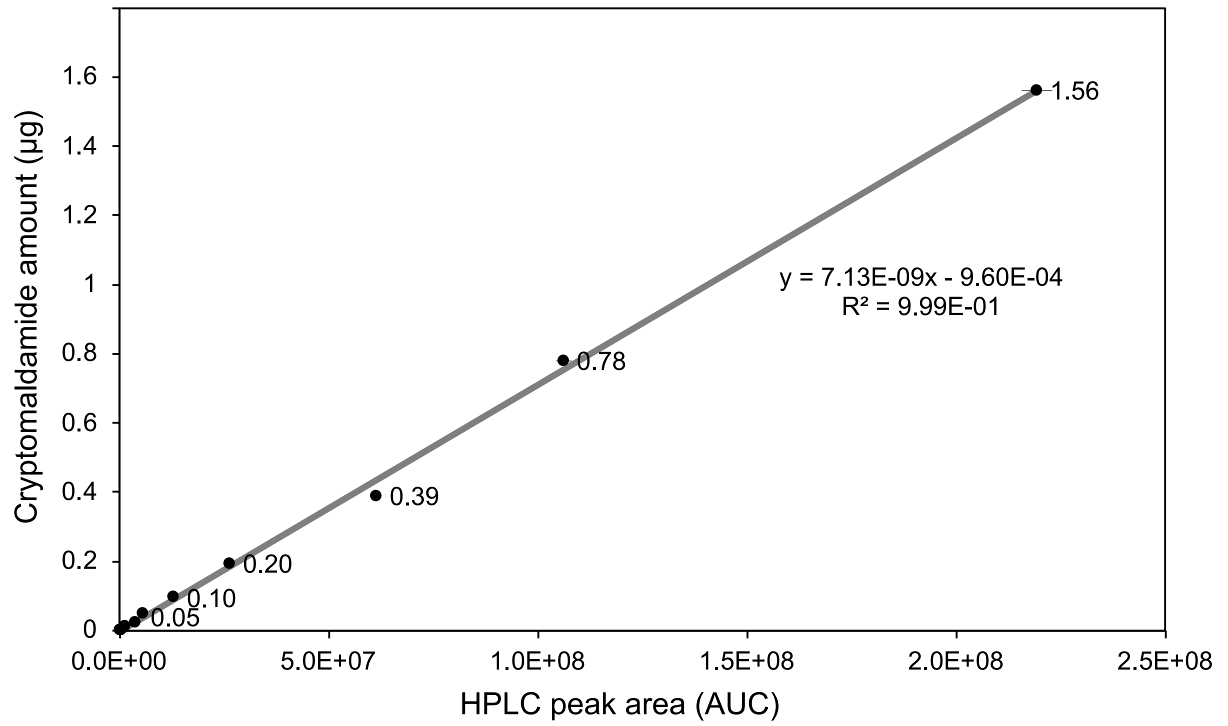


**Figure S4.** LC-MS generated standard curve plotting detector response area under the curve (AUC) for the $(M+H)^+$ peak at *m/z* 400 versus the amount of cryptomaldamide standard injected. Amounts of cryptomaldamide injected (µg) were 0.001, 0.003, 0.006, 0.012, 0.024, 0.049, 0.098, 0.195, 0.391, 0.781, 1.5625. The trend line and the equation were determined in Excel.

### TABLES

**Table S1. Sequencing results listing mutations for 6 plasmids carrying the cryptomaldamide gene cluster**

| Clone | *cpmA* | *cpmA*-*cpmB* intergenic | *cpmB* | *cpmB*-transposase intergenic | transposase |
|---|---|---|---|---|---|
| CR03 | | +1243/-2695 (G→A) | R1199R (CGG→CGA)<br>E2086G (GAA→GGA)<br>A2683S (GCC→TCC)<br>L4494F (TTG→TTT)<br>S4610L (TCG→TTG) | | P139P (CCG→CCA) |
| CR15 | | +1243/-2695 (G→A)<br>+2148/-1790 (T→A) | A1522A (GCT→GCC)<br>G5049C (GGC→TGC) | | |
| CR31 | | +1243/-2695 (G→A) | P220P (CCA→CCG)<br>C1250C (TGC→TGT)<br>L2050I (CTA→ATA) | | |
| CR33 | | +1243/-2695 (G→A) | Q88K (CAG→AAG)<br>P3544T (CCA→ACA) | | |
| CR84 | E176G (GAG→GGG)<br>F204Y (TTT→TAT) | +1243/-2695 (G→A)<br>+1441/-2497 (T→C) | P3850P (CCC→CCT)<br>R4767H (CGC→CAC)<br>L6130I (CTC→ATC) | +243/-1552 (C→A) | |
| CR92 | | +1243/-2695 (G→A) | | | |

**Table S2. $^1$H Proton NMR Comparison of Cryptomaldamide produced in *M. producens* and *Anabaena***

| *Position* | *Reported ($D_2O$ 800MHz)*[1] | *Observed (DMSO d-6 500MHz)* |
|---|---|---|
| *1* | | *7.78 s* |
| *3* | | *8.03 s* |
| *4* | | *8.09 s* |
| *5* | *4.23 dd* | *4.18 dd* |
| *6* | *3.78 dd* | *3.05 dd* |
| *11* | | *8.3 d* |
| *12* | *4.45 d* | *4.43 m* |
| *14* | *1.95 m* | *1.92 m* |
| *15 16* | *0.78 d* | *0.77 d* |
| *20* | *2.95 s* | *2.87 s* |
| *19* | *4.72 dd* | *4.82 dd* |
| *21* | *1.85 m* | *1.80 m* |
| *22 23* | *0.65 d* | *0.68 m* |
| *24* | *6.21 dq* | *6.52 d* |
| *26* | *1.68 d* | *1.70 s* |

**Table S3. Plasmids**

| Plasmid name | Description | | Source |
|---|---|---|---|
| pCAP03-acc(3)-IV | Expression vector for yeast and bacteria | Km | [2] |
| pCVD002 | $Sp^R$, $Sm^R$ gene carried on a CYANO-VECTOR device | Ap, Sp, Sm | [3] |
| pCVD003 | $Km^R$ gene carried on a CYANO-VECTOR donor plasmid | Ap, Km | [3] |
| pCVD015 | Counter selectable *ccdB*-based cloning cassette carried on a CYANO-VECTOR donor plasmid | Ap | [3] |
| pCVD022 | *S. elongatus* NS2 carried on a CYANO-VECTOR donor plasmid | Ap | [3] |
| pCV0095 | Plasmid for chromosomal integration at *S. elongatus* NS3 carrying *B. subtilis* sfp-PPTase. | Cm | [4] |
| pAM5273 | pCVD022 with the yeast element ARSH/CEN6 element and *trp1* | Ap | This study |
| pAM5276 | *ura3* gene under the *Schizosaccharomyces pombe* Padh1 promoter carried on a CYANO-VECTOR donor plasmid | Ap | This study |
| pAM5277 | pAM5576 carrying the point mutation T189C in *ura3* | Ap | This study |
| pAM5372 (pJR01) | pANS-based replicative expression vector for *S. elongatus.* The vector carries a counter selectable *ccdB*-based cloning cassette downstream of the PconII promoter. | Sp, Sm | [4] |
| pAM5406 | CYANO-VECTOR donor plasmid carrying an RSF1010 replicon with substitution mutation Y25F in the *mobA* gene that inactivates MobA's nicking activity and the addition of RK2-bom site and pUC19 origin of replication for efficient conjugal transfer and high copy number of the plasmid in *E. coli.* | Ap | [5] |
| pAM5544 | Plasmid for chromosomal integration at *S. elongatus* NS2 carrying a $Nt^R$ gene | Nt | [6] |
| pAM5564 | pAM5571 carrying the Cryptomaldamide BGC, clone 92 | Km | This study |
| pAM5565 | pAM5572 with gRNA spacer sequence targeting $Nt^R$ gene integrated into *Anabaena* AMC2556 chromosome. | Sp, Sm | This study |
| pAM5569 | pER015 carrying *S. elongatus* NS2 homology region flanking $Nt^R$ gene | Nt | This study |
| pAM5571 | Yeast capture vector for recombination in *S. elongatus* at NS2 | Km | This study |
| pAM5572 | Improved RSF1010-based broad host range plasmid (using pAM5406 replicon) for genome editing using Cpf1/CRISPR technology. | Sp, Sm | This study |
| pAM5600 | Cpf1/CRISPR module (SalI-PstI fragment) from pSL2680 carried on a CYANO-VECTOR donor plasmid. | Ap | This study |
| pAM5601 | pAM5372 carrying *Anabaena* sfp-type PPTase *hetI* (all5359) | Sp, Sm | This study |
| pER015 | Plasmid for chromosomal integration at *Anabaena* NS2 carrying a counter selectable *ccdB*-based cloning cassette and the *sacB* gene for positive selection of double recombinant clones on sucrose. | Km | [3] |
| pSL2680 | RSF1010-based broad host range plasmid for genome editing using Cpf1/CRISPR technology (Addgene no. 85581). The plasmid comprises *Francisella novicida cpf1* gene, a CRISPR array with a lacZ$\alpha$ fragment flanked with AarI restriction sites to facilitate the cloning of a guide RNA spacer sequence, and a SalI-KpnI site for the cloning of a homologous repair template. | Km | [7] |

**Table S4. Oligonucleotides**

| Oligonucleotide name | Sequence (5'-3') | Template and comments |
|---|---|---|
| **CYANO-VECTOR TAR cloning modules** | | |
| S7942NS-Yeast-F | ttttgctggccttttgctcacataaaataaacaaataggggttccgc | Yeast plasmid replication and selection elements |
| S7942NS-Yeast-R | ggggataacgcaggaaagaataggttcacgtagtgggccatc | |
| Padh_u38F | cggccaataacccagggatttaatagctgcgccgatggtttctac | Yeast cloning destination cassette |
| ura3_D810R | tgccggggagctccttcatttggtaccttagttttgctggccgc | |
| ura3-t189c-F | gtttactaaaaacacatgtggacatcttgactg | To quick change the EcoRV site in the *ura3* gene of pAM5276 |
| ura3-t189c-R | aatcagtcaagatgtccacatgtgtttttag | |
| **Screening for the BGC** | | |
| aphI_528R | ttcaacaggccagccattacgc | Plasmid – BGC junction |
| crypt_215R | ggcagggagcaccaagaaactg | |
| crypto_2015F | actctatcttggccctcacttctcttg | BGC – *cpmB* gene |
| crypto_2788R | tgttgttttgttggttgtttggtttg | |
| crypt_27823F | ggggcattttattgtcttggaagc | BGC – plasmid junction |
| S7942NS2LA_274R | gacctagatgaggaagcatgagcg | |
| **Capture of the BGC** | | |
| P100F | ggtttgacgcctcccatggtataaatagtggctcgagctcttaacg ctctatcccttatc | Cryptomaldamide BGC (1 – 10155 bp) with 30 bp plasmid recombination sequence |
| P100R | ggaaactcaaccgtcgtctg | |
| P101F | cgacagttatcaatggttggtag | Cryptomaldamide BGC (10024 – 19823 bp) |
| P101R | ctttaggtttaagaggaatgcc | |
| P102F | gattgggcaacagatgtcctg | Cryptomaldamide BGC (19655 – 28095 bp) with 30 bp plasmid recombination sequence |
| P102R | cagcacgttccttatatgtagctttcgacatcgcctgaagtcagaca ttttatcagaggg | |
| **RT-qPCR** | | |
| cpmA_103FQ | gcaacagttcctccggattc | *cpmA* (1, 103 – 262 bp), for RT-qPCR |
| cpmA_262RQ | ccggttgtcgaatcttgacc | |
| cpmA_657FQ | gcgccgtcacttaggagata | *cpmA* (2, 657 – 754 bp), for RT-qPCR |
| cpmA_754RQ | cgggagccattggcataaa | |
| cpmB_4215FQ | aaccctcgggctgtttatca | *cpmB* (1, 4215 – 4355 bp), for RT-qPCR |
| cpmB_4355RQ | tgcacgtctaccaaggaact | |
| cpmB_8269FQ | atcattcttgcccggaaacg | *cpmB* (2, 8269 – 10808 bp), for RT-qPCR |
| cpmB_10808RQ | acgggagtcacatcccattt | |
| cpmB_15710FQ | acccattggctagcagaact | *cpmB* (3, 15710 – 15835 bp), for RT-qPCR |
| cpmB_15835RQ | cttgaagcacctcctgcatc | |
| 0649_359F | ttgctctagaacgcattcgc | *secA* (359 – 507 bp), for RT-qPCR |
| 0649_507R | ctgcaccatcttgtccttgg | |
| 0289_144F | gaagctcgacaaggtttccc | *rpoD* (144 – 325 bp), for RT-qPCR |
| 0289_325R | ccgtcttcatctcggcaatc | |
| **HetI expression plasmid** | | |
| pAM5372_A7120_hetI_2F | aaataaaggaggtcttaagatggaaccagggcaagttaaatttg | *hetI* gene (all5359 in *Anabaena* gDNA) |
| pAM5372_A7120_hetI_583R | caggatggccttctcctgcatcaataatgccagaattttggctgc | |
| **CYANO-VECTOR CRISPR module** | | |
| pCVD015_1684F | taccgtcgacatcgggggcccccggggggacgtccaggggata acgcaggaaagaac | pCVD015 backbone with SalI and CYANO-VECTOR GC-linker |
| pCVD015_3464R | atgcctgcagatccggcggcggcggcgggacgtcgcggaacc cctatttgtttatttttc | |
| **A7120NS2** | | |
| A7120NS2xS7942NS2LAF | atatcaggacatatccaccaggccacgatttgagggacgaatctt c | *S. elongatus* NS2 homology sequence |
| A7120NS2xS7942NS2RAR | ctcactagggtaagcaaagctatcaggattaatgaaacggacgc cc | |
| **CRISPR gRNA** | | |
| NT_U248F | agatgacagcttatcatcgaatta | gRNA template for the nourseothricin resistance gene in AMC2556 |
| NT_U228R | agactaattcgatgataagctgtc | |

**Table S5. Strains**

| Strain | Description | Resistance | Source |
|---|---|---|---|
| **Escherichia coli** | | | |
| NEB Stable | Cloning strain | Tc, Sm | NEB |
| DH5a | Cloning strain | | |
| AM1359 | Conjugal strain, *E. coli* DH10B carrying pRL623 and pRL443 helper and conjugal plasmid, respectively. | Ap, Tc, Cm, Sm | [8] |
| **Saccharomyces cerevisiae** | | | |
| *S. cerevisiae* VL6-48N | TAR cloning strain | | [9] |
| **Cyanobacteria** | | | |
| AMC0001 | *Anabaena* (*Nostoc*) sp. PCC 7120 WT | | Laboratory collection |
| AMC2302 | *S. elongatus* PCC 7942 WT cured of its small plasmid pANS | | [10] |
| AMC2566 | AMC2302 carrying *B. subtilis* sfp PPTase driven by PconII-riboswitch-F at NS3 | Cm | This study |
| AMC2556 | *Anabaena* sp. PCC 7120, engineered to contain *S. elongatus* neutral site 2 inserted into the all1697 gene | Nt | This study |
| AMC2560 | AMC2556 carrying cryptomaldamide pathway – clone 1 | Nt, Nm | This study |
| AMC2561 | AMC2556 carrying cryptomaldamide pathway – clone 2 | Nt, Nm | This study |
| AMC2562 | AMC2556 carrying cryptomaldamide pathway – clone 3 | Nt, Nm | This study |
| AMC2563 | AMC2560 conjugated with pAM5565 to force segregation – clone 1 | Nm | This study |
| AMC2564 | AMC2560 conjugated with pAM5565 to force segregation – clone 2 | Nm | This study |
| AMC2565 | AMC2560 conjugated with pAM5565 to force segregation – clone 3 | Nm | This study |
| *Moorena* (*Moorea*) *producens* strain JHB | Marine cyanobacteria, native host of cryptomaldamide | | Laboratory collection |
